# SARS-CoV-2 sequence typing, evolution and signatures of selection using CoVa, a Python-based command-line utility

**DOI:** 10.1101/2020.06.09.082834

**Authors:** Farhan Ali, Mohak Sharda, Aswin Sai Narain Seshasayee

**Author notes:** To whom correspondence should be addressed. Farhan Ali; Aswin Seshasayee.

## Abstract

The current global pandemic COVID-19, caused by SARS-CoV-2, has resulted in millions of infections worldwide in a few months. Global efforts to tackle this situation have produced a tremendous body of genomic data, which can be used for tracing transmission routes, characterization of isolates, and monitoring variants with potential for unusual virulence. Several groups have analyzed these genomes using different approaches. However, as new data become available, the research community needs a pipeline to perform a set of routine analyses, that can quickly incorporate new genome sequences and update the analysis reports. We developed a programmatic tool, CoVa, with this objective. It is a fast, accurate and user-friendly utility to perform a variety of genome analyses on hundreds of SARS-CoV-2 sequences. Using CoVa, we define a modified sequence typing nomenclature and identify sites under positive selection. Further analysis identified some peptides and sites showing geographical patterns of selection. Specifically, we show differences in sequence type distribution between sequences from India and those from the rest of the world. We also show that several sites show signatures of positive selection uniquely in sequences from India. Preliminary evolutionary analysis, using features that will be incorporated into CoVa in the near future, show a mutation rate of 7.4 × 10^−4^ substitutions/site/year, confirm a temporal signal with a November 2019 origin of SARS-CoV-2, and a heterogeneity in the geographical distribution of Indian samples.

## Introduction

Being one of the biggest public health crises that the world has faced in the 21st Century, COVID-19 needs no introduction. The causal agent of this pandemic is a novel Coronavirus, a group of positive strand RNA viruses which also includes SARS and MERS, and is identified as SARS-CoV-2 (1). In a short span of 5 months, the virus has expanded globally and has accumulated a large number of variants (2), thus requiring the continued monitoring of variation in its population. With a large body of genomic data publicly available, preprint servers have seen a deluge of variant analysis reports. However, these reports typically differ in their methodology and the extent of analysis. As more and more genomes are being sequenced, we require a rapid and routine tracking of variation in the population as well as genomic characterization of these new isolates.

To serve this purpose, we developed CoVa, a pipeline for Coronavirus variant analysis. CoVa is a fast, light-weight and user-friendly command-line tool, especially geared to analyse hundreds to a thousand of SARS-CoV-2 genomes. CoVa not only calls variants but bundles several routine analyses required to trace the progress of this disease in genomic terms. This includes estimation of sequence diversity, type identification, phylogeny and selection analysis, with identification of sites potentially undergoing positive selection. We use CoVa to describe sequence diversity and signatures of selection in SARS-CoV-2 genomes with special emphasis on sequences from India.

## Methods

### Implementation

CoVa is implemented as a python library meant to be run as a command-line tool. The only required input is a multi-FASTA file of assembled whole-genome sequences. CoVa first builds a whole-genome multiple sequence alignment (wgMSA) which serves as the starting material for all downstream analyses. NCBI Refseq accession NC_045512 is used as the variant calling reference in the pipeline. Therefore, this genome is required to be included in the input file. Since several analyses require identification of “reference” sites, CoVa includes a command to reduce wgMSA to sites present in the reference genome (rMSA). Duplicate sequences are removed at this stage (generating uMSA) to speed up downstream analyses and to avoid polytomies in phylogeny. The entire set of commands along with their input and output transfers have been depicted as a directed acyclic graph in **Figure 1**. Cova has been implemented in a way that allows for execution of individual commands or a combination thereof. It is also possible to run the entire pipeline with a single command.

**Figure 1:**
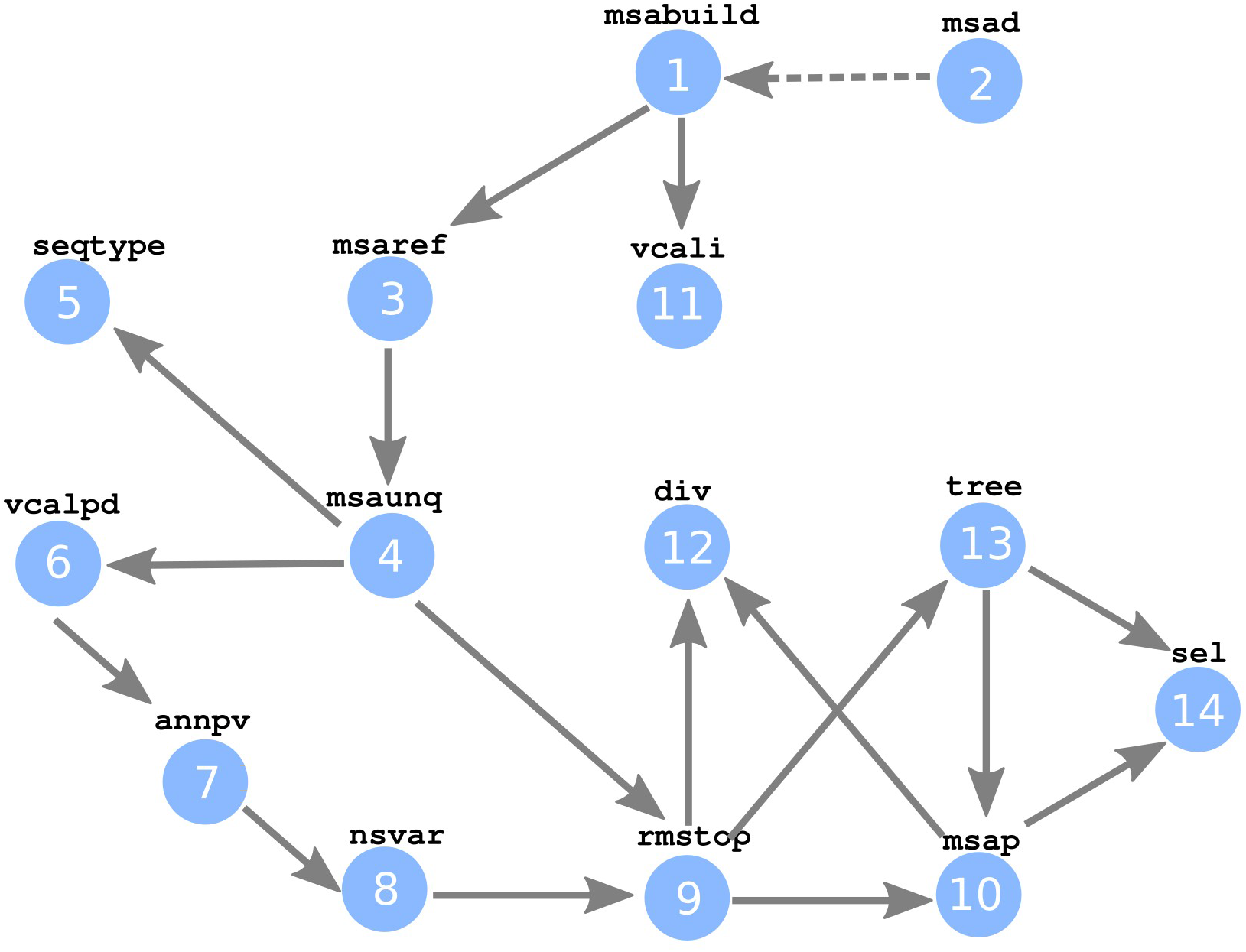
Cova’s workflow. Schematic of CoVa’s workflow with all its sub-commands and their input/output connections.

CoVa splits into multiple programs following the generation of uMSA, viz., variant calling, diversity computation, phylogeny and selection analysis. Selection analysis incorporates estimation of non-synonymous and synonymous rates for each region, and identification of individual sites, if any, under positive selection. Sequences with nonsense mutations throw an error in the selection analysis. Therefore, these sequences are removed from the alignment, generating sfMSA. Selection analysis also requires extraction of protein/peptide-encoding nucleotide regions of MSA (pnMSA), along with the phylogenetic tree, both of which also need to be generated from sfMSA. The pnMSAs are also used to compute nucleotide diversity of these regions, along with that of the whole-genome. CoVa also includes an option to calculate nucleotide diversity with a window sliding over the entire genome. Besides its core functionality, CoVa packages several functions to preprocess sequences, extract their metadata and for plotting phylogenies.

CoVa employs several popular programs for individual jobs, viz., MAFFT (3) to build wgMSA, FastTree2 (4) for phylogeny, and FUBAR (5) program of Hyphy to identify sites under positive selection. These programs were selected for their speed and accuracy. CoVa runs these commands with such options that optimize for speed without a heavy compromise on the accuracy. For example, CoVa uses a maximum of 5 iterative refinements in MAFFT, as opposed to MAFFT’s speed-default (2) and the accuracy default (1000). Given that most input genomes are expected to be highly similar and MAFFT achieves most of the accuracy in the initial refinements, a limit of 5 allows CoVa to be fast and accurate, as well as consistent in its runtime. CoVa switches to MAFFT’s speed-default if the number of input sequences exceeds 1000. Similarly, CoVa limits split-support computation in FastTree to 100 runs for both speed and memory optimization without compromising on accuracy.

One of the key advantages of using MAFFT in CoVa is its ability to quickly incorporate new sequences to an existing MSA (6). CoVa provides this feature explicitly for a prompt integration of new data and updation of existing analysis results. With this option, labs can compare and integrate their local genomic data with the sequences already available in the public repository. To facilitate this, an MSA comprising 825 sequences from GenBank is available on CoVa’s github page.

### Estimation of Sequence diversity

The extent of variation in the population is estimated in terms of nucleotide diversity, which is the average pairwise difference per-unit length. CoVa can compute the whole-genome nucleotide diversity of an MSA of thousands of sequences in seconds. It achieves this by splitting the calculation over individual sites, which reduces the time complexity to ****O(NL)**** as opposed to ****O(N^2^)****, where *L* is the number of sites and *N* is the number of sequences.

Specifically, we calculate nucleotide diversity **π** as follows:

Let ****B**** be an ordered set of alphabets, *e.g.*, {A,C,G,T} with size *K*. *f*_i_ represents the number of occurrences of *B*_*i*_ for a single site in the MSA. Then,

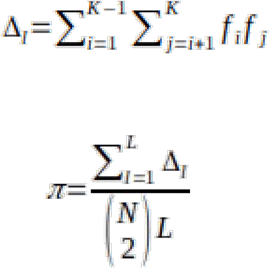

where, *Δ*_*l*_ = total pairwise base difference for site *l.*

CoVa also estimates diversity of individual peptide-encoding regions. Optionally, following (7), the entire genome can be scanned with a sliding window diversity calculator to identify segments of high diversity. This is a convenient way to quickly observe high-frequency variants and the affected genomic loci. Most of these hotspots also align with the positions used in (8) to type SARS-CoV-2 sequences, and can be instrumental in future for an automated sequence typing scheme for this virus.

### Sequence typing with CoVa

Genome sequences of SARS-CoV-2 are typed in CoVa following the barcoding scheme suggested in (8). This involves extracting a nucleotide sequence from 10 positions spread across the genome. The positions are listed in **Table 1**, along with the affected peptide and its major amino acid substitution. Every variant of this 10-nucleotide long sequence tag is considered a distinct sequence type. Sequence types are indexed in the order of their earliest collection date, *i.e.*, at least one sequence of type ST-*i* was observed before any sequence of type ST-*i+1*.

**Table 1.** 10 nucleotide positions used for sequence typing in CoVa, along with their major mutation.

| SN | Position | Peptide | Mutation |
| --- | --- | --- | --- |
| 1 | 1397 | nsp2 | V198I |
| 2 | 1440 | nsp2 | G212D |
| 3 | 2891 | nsp3 | A58T |
| 4 | 3037 | nsp3 | F106F |
| 5 | 8782 | nsp4 | S76S |
| 6 | 14408 | RDRp | P323L |
| 7 | 23403 | S | D614G |
| 8 | 26144 | ORF3a | G251V |
| 9 | 28144 | ORF8 | L84S |
| 10 | 28688 | N | L139L |

### Data Sources

CoVa was initially validated on whole genome sequences of 825 SARS-CoV2 isolates from GenBank. For the analysis described in this study, sequences were acquired from GISAID (accessed May 21). Only complete and high coverage genomes were downloaded. All of the 301 Indian genomes were analyzed. To select a sample from over 30,000 sequences available from other countries, the GISAID browser option of selecting sequences with known patient’s status was used. This limited the “global” dataset (countries excluding India) to 1356 genomes. This data was further processed to exclude all genomes with more than 1% ambiguous characters, retaining 248 Indian and 1262 global genomes. Since CoVa removes duplicate genomes from MSA, our final dataset had 244 Indian and 1129 global sequences.

Information on countries’ population was collected from Worldometer (https://www.worldometers.info/world-population/population-by-country/) and on countries’ latitude was collected from Google’s public dataset (https://developers.google.com/public-data/docs/canonical/countries_csv).

### Evolution of SARS-CoV-2

Two multiple sequence alignments built using - 1) only Indian samples and 2) samples across the globe (excluding Indian samples) were merged together as a single multiple sequence alignment (MSA) using the mafft --merge option (MAFFT reference). The effect of sample over-representation bias was reduced by including one sample per isolation date for a given geographical location. Therefore, the final alignment used for phylogenetic tree reconstruction consisted of 413 SARS-CoV2 whole genome sequences (WGS), including the reference genome. The conserved regions relevant for phylogenetic inference were extracted from the MSA using BMGE v1.12, with -DNA option. Using IQTREE v1.6.5 (9), a maximum likelihood (ML) based tree was built with GTR+F+I nucleotide substitution model. ModelFinder (10) with -m MF option was used to choose this model compared against 285 other model combinations based on the least Bayesian Information Criterion (BIC) score. This pipeline was created using python 2.7.

This procedure for generating trees, which is more accurate (11) but substantially slower than FastTree, is not part of CoVa at present in light of CoVa’s preference for faster methods. However, the accuracy of this procedure could be critical to more sophisticated evolutionary analysis including ancestral reconstruction, and will be considered for a future update of CoVa.

To examine the relationship between time (temporal signal) and genetic divergence we used the ML tree produced above which assumes a non-molecular clock model. We used TempEst (12) to find the root of the tree such that it optimised for the temporal signal by trying all possible roots and chose the one that minimised the mean of the square of the residuals. This also allowed us to estimate the rate of evolution of SARS-CoV2, reported as substitutions per site per year.

### Clustering of sites under selection

Amino acid sites were clustered based on their probability of positive selection across 13 countries. A site with its probability of non-synonymous rate (**β**) being greater than synonymous rate (**α**), *P*_β>α_ > 0.7 was considered to be under positive selection. This threshold was based on the distribution of these values across sites. The probabilities across countries were transformed into bits; 1 for positive selection and 0 for not. Binary distance (proportion of bits in which only one is 1, out of all the bits in which at least one is 1) was calculated for all pairs of sites. Indices of site pairs with zero distance were collected and clusters were identified based on overlapping indices.

## Results

### CoVa’s Performance evaluation

We tested CoVa’s speed on multiple pilot datasets, of different sizes *viz.*, small (30), medium(106) and large(826). Starting from the un-aligned sequences, CoVa finished the entire analysis in approx. 2, 6 and 50 minutes for small, medium and large datasets respectively, using only 4 CPUs. To test its accuracy on variant calling, we had initially compiled a set of 21 mutations described in the SARS-CoV-2’s literature from the preprint server BioRxiv (till May 03) (**Sup. table 1**). CoVa correctly identified 19 of these mutations. The two mutations where CoVa’s identification differed from that of the original authors were originally displaced by 1 residue. Therefore, the variant calling feature of CoVa showed 100% accuracy. This also validates the MSA-building procedure used in CoVa, as the accuracy of variant calling would be directly affected by the accuracy of the MSA. As for the phylogeny and selection analysis, CoVa can be expected to be as accurate as the external programs it invokes for these purposes. It is worth noting in this regard that even though the Spike protein appears overall conserved, FUBAR identified multiple sites potentially under positive selection, corroborating previous reports (13,14). On the other hand, it supported the hypothesis in (15) of positive selection on ORF8 but not on ORF3a, using the large dataset. However, ORF3a was predicted to be under positive selection on the medium dataset. This likely reflects the fact that the selection on ORF3a was not pervasive, and therefore not amenable to detection through FUBAR (we address this specifically below).

All analysis described below are based on the larger dataset described in Methods.

### Sequence types of SARS-CoV2

We have used the ten nucleotide genetic barcoding scheme suggested by Guan *et al*, 2020 (8) to identify clades in SARS-CoV2 phylogeny. We have opted for a different nomenclature than the one used in (8) or the one adopted by GISAID. In CoVa, genomes are identified and categorized into “sequence types” (defined by the barcoding scheme), as opposed to “clades”. Moreover, as opposed to naming clades after a specific amino acid substitution, these types were named in the order of their first appearance. For example, the first sequence to be typed ST1 must have been observed before any sequence of type ST2. This scheme has the advantage that the new types are naturally accommodated in the nomenclature as they emerge and the standing variation can be captured at a finer resolution as compared to a hard-coded clade nomenclature.

Our typing system identified 18 sequence types in the rest-of-the-world (referred to as “global” in the rest of this paper) dataset, 5 of which had more than 1% representative sequences. We looked for the distribution of these 5 types (ST1, 3, 4, 5, 10) in the Indian dataset (**Figure 2A)**. We found ST1 and ST4 to be overrepresented while ST3 and ST10 to be underrepresented among the Indian genomes (*P*_Binom_ < 0.01, adjusted for multiple testing using “Holm-Bonferroni correction”). Other types only amounted to 2 %. 3 Indian sequences could not be typed due to the presence of ambiguous characters at the barcoding positions. Sequence type information of all the analyzed genomes is available from **Sup. table 2**.

**Figure 2:**
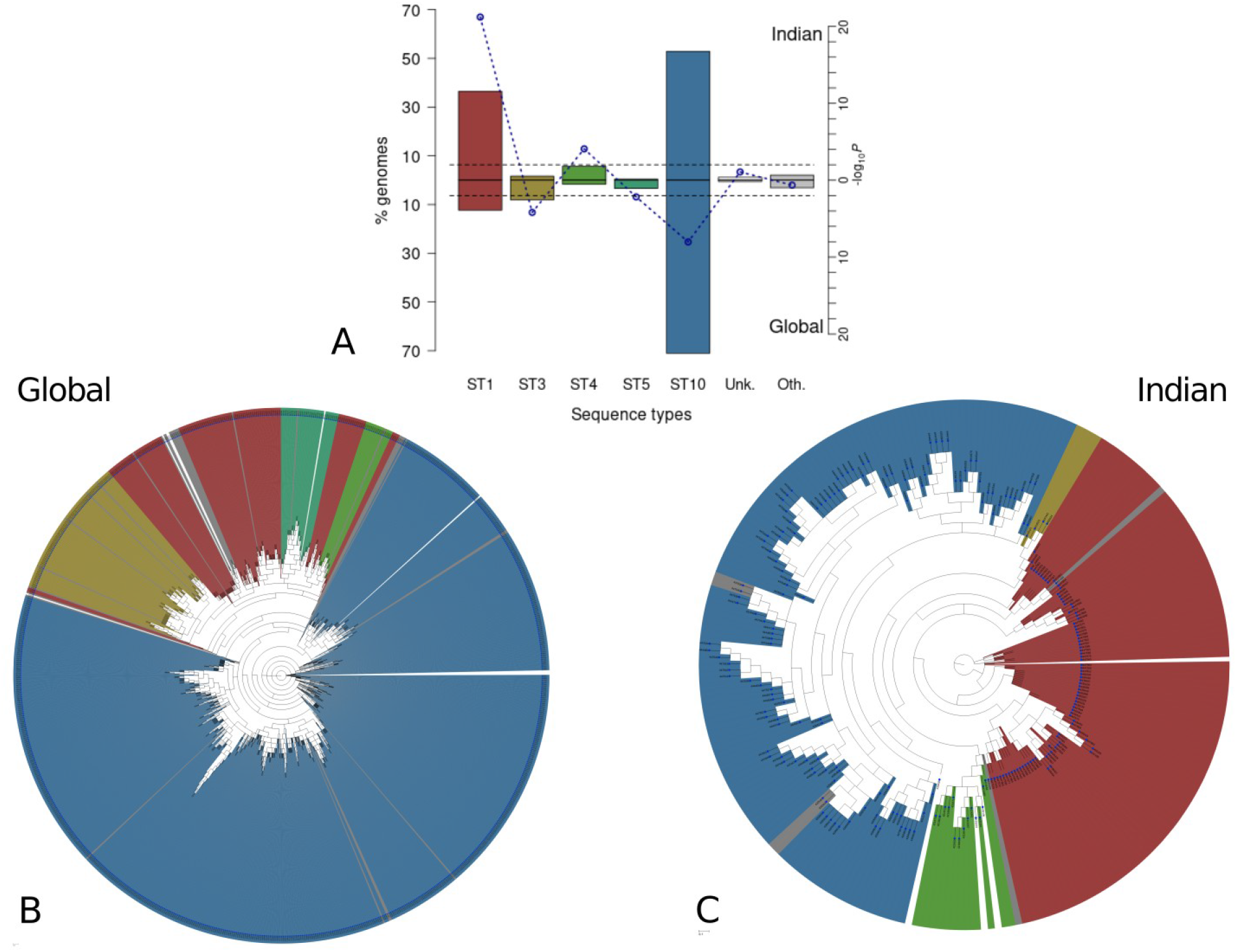
Sequence types of SARS-CoV-2. (A) Distribution of sequence types of SARS-CoV-2 genomes sequenced in India, in comparison with their global distribution. Whole genome phylogeny of global (B) dataset and Indian (C) dataset, showing phylogenetic clusters of these sequence types.

We built separate whole-genome phylogenies for the Indian and the global datasets using FastTree2, as implemented in CoVa. The barcoding scheme used in CoVa successfully clustered sequences into distinct clades (**Figure 2B,C**). However, several sectors of type ST1 can be seen in the global phylogeny (**Figure 2B**). This is not due to any limitation on our choice of phylogeny method and software, as multiple variants on CoVa’s implementation of FastTree2 as well as other softwares like MEGAX (16) (used in GISAID), and FastME2 (17) were tried and they did not improve upon these results (not included in this study). Rather, it highlights the fact that a 10 nucleotide barcode cannot capture the entire emergent diversity of SARS-CoV2 genomes. This is where our choice of using “ sequence types” over “clades” becomes significant as types do not have to form mutually exclusive phylogenetic clusters. Each separate sector of ST1 was essentially a different clade which could not be typed distinctively by the present barcode. As an example to this effect, a distinct clade in India was reported in (18). We noticed that CoVa categorized this cluster as ST1. In fact, 85 of 88 ST1 sequences from India belonged to this cluster. Since none of the 4 positions used to define this cluster (6312, 13730, 23929, 28311) were included in the genetic barcode, it did not receive a distinct sequence type. However, 4 sequences with the same mutations were of type ST10 (3) and ST17 (1), possibly as a result of recombination.

### Evolution of SARS-CoV2

Our analysis - using the dated phylogeny, which is not yet incorporated within CoVa (see Methods, **Sup. table 3**) - suggests that SARS-CoV2 is a measurably evolving population, with a sufficient divergence accumulation over our sampling time range (R^2^=0.25, correlation coefficient, r = 0.5, Figure 3A). We obtained an evolutionary rate of 7.4 × 10^−4^ substitutions/site/year and the time to the most recent common ancestor (TMRCA) of late November 2019 (**Figure 3A**), the latter being consistent with recent studies (19–21). The evolutionary rate estimation, however, is lower than what has been reported previously in coronaviruses i.e of the order of 10^−3^ substitutions/site/year (22–24), but is similar to that described by others for SARS-CoV-2 (25) (https://bit.ly/2BRvEwj). Our estimate is also similar to those reported for SARS and MERS coronaviruses (22,26) and is a third of the estimates reported for influenza B (27). As noted before (28), root-to-tip regression, employed in this study, often produces lower estimates than those from Bayesian methods. For our dataset, whether this is a technical artifact remains to be tested.

**Figure 3:**
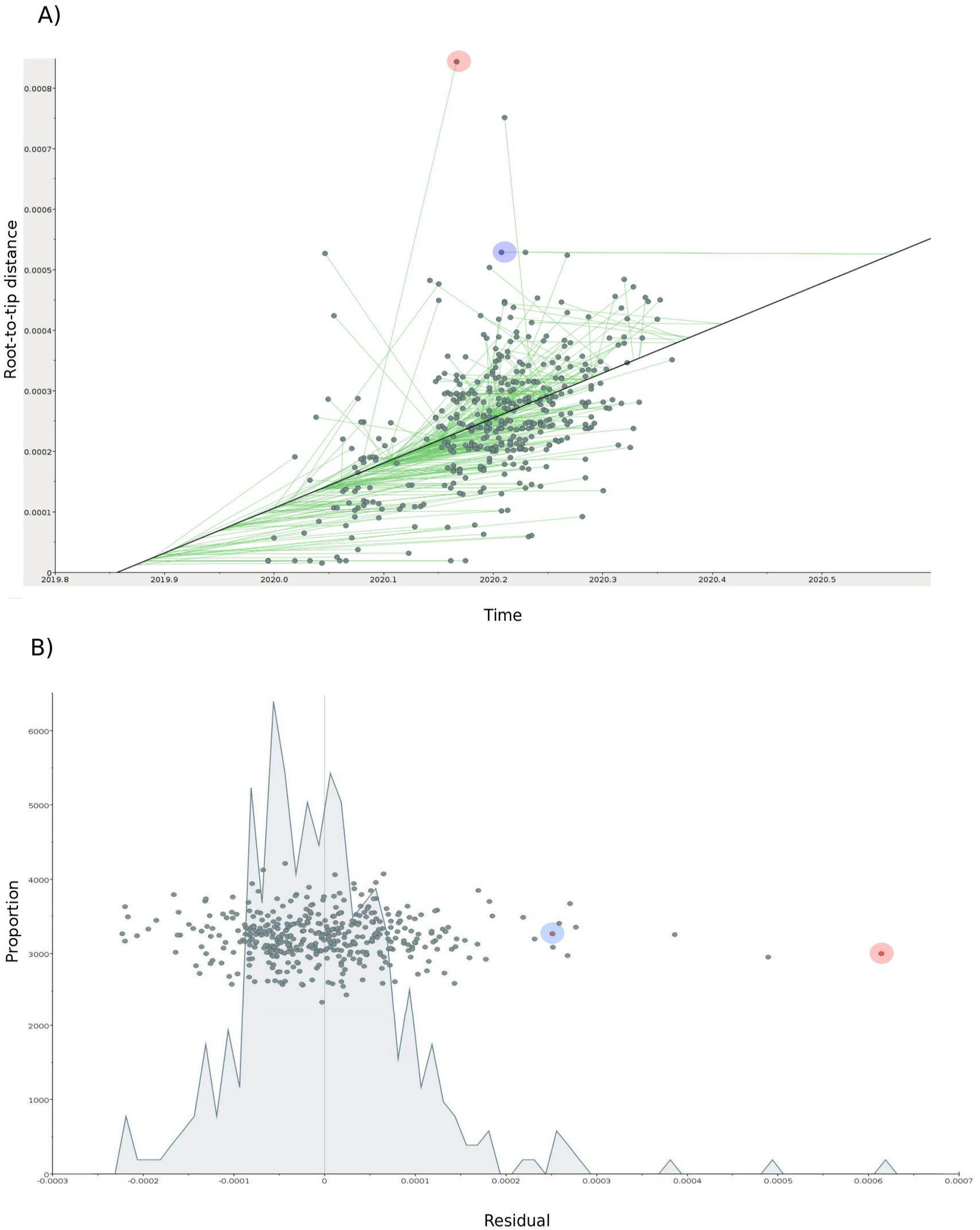
Temporal signal analysis of SARS-CoV-2. A) Scatter plot showing the relationship between the root-to-tip distances (genetic divergence) and time (calculated as sampling times). The best fit line represents the minimum residual mean squared test statistic. Green lines represent the ancestral traces for each sample. (B) Scatter plot showing the relationship between the residuals for each sample and their proportion in the dataset.

Ancestor tracing of sequences like the one isolated from Taiwan on 17th March 2020 (Accession number: 422415) suggested that the sequence might be more recent than the date it has been given (Figure 3A and 3B, red circle). Another outlier sequence type like the one isolated from France on 2nd March 2020 (Accession number: 414628) was substantially more divergent from its ancestor than expected given a linear relationship between time of divergence and genetic diversity (Figure 3A and 3B, blue circle). These observations can be explained in a number of ways. For example, the former could be a result of recombination from a recent virus of the supposed date. Similarly, the latter could be because of sequencing errors, recombination etc. One more explanation that could give rise to these “outliers” is the difference in the rate of evolution among different lineages i.e a relaxed molecular clock as opposed to a strict molecular clock assumed in this analysis. In this direction, a recent study of sequences from India has suggested that different lineages are evolving at different rates (18). However, since the outbreak recently emerged, the number of substitutions is still small. Therefore, the confounding factors described above are a more likely explanation of the large residual values (**Figure 3B**) observed for our samples than a molecular clock variant. Only an extensive Bayesian analysis based study comparing different variants of molecular clock models would shed more light in this direction.

The dated phylogeny obtained after applying the best fitting root (see Methods) shows the diversification of SARS-CoV2 along two distinct lineages with respect to the root - one temporally closer to the root than the other (**Figure 4**). The former lineage contains thirty samples (88%) submitted from China, including the reference genome reported in late December 2019. Samples from India are distributed across the two main lineages identified, 43% samples belonging to the lineage temporally closer to the root. Indian samples cluster with those belonging to different geographical locations. A further ancestral reconstruction analysis would shed light on the most likely origins of this heterogeneity identified among the Indian samples.

**Figure 4:**
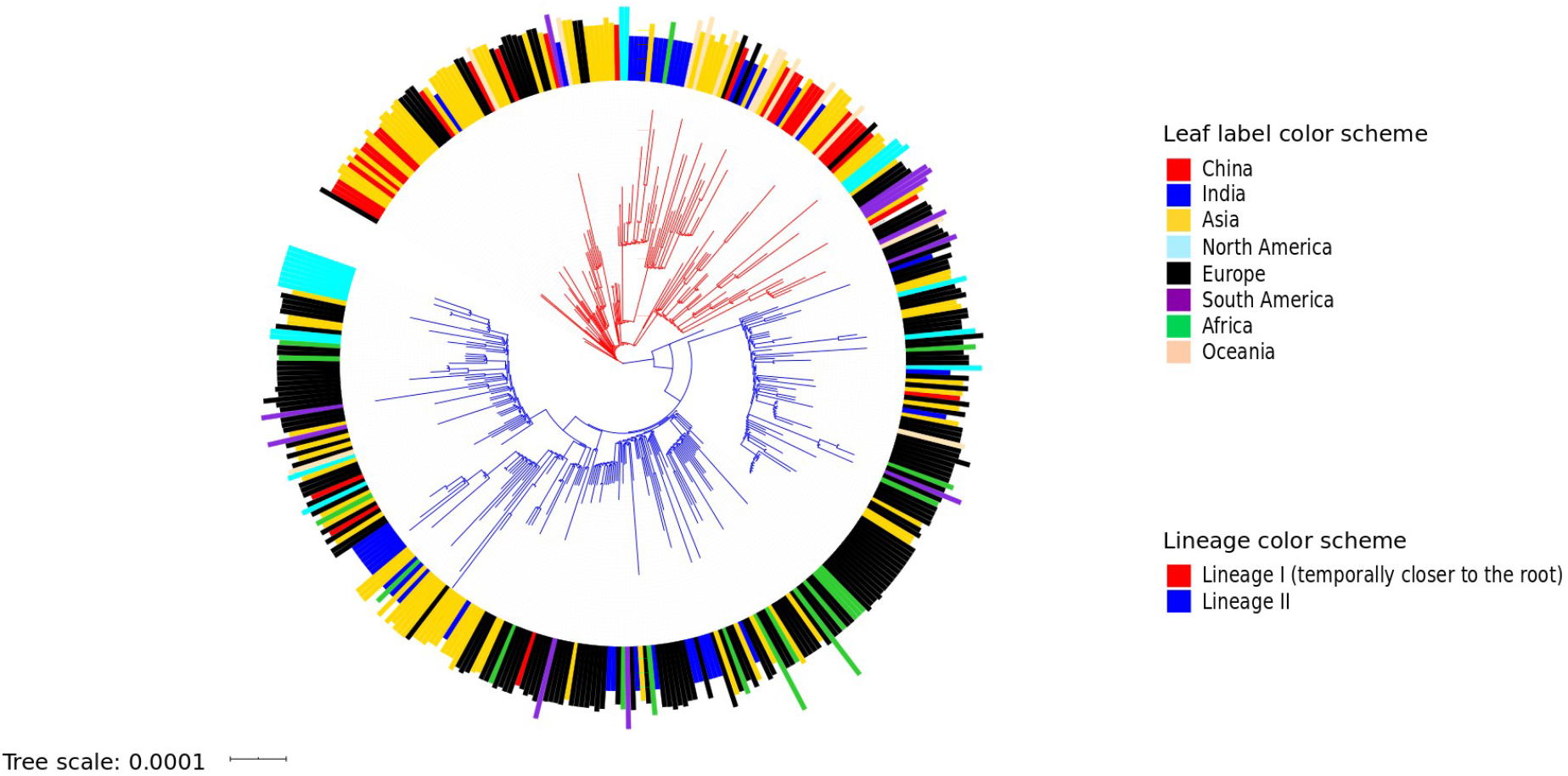
Maximum-likelihood based dated tree with best fit root. A dated Maximum likelihood based phylogenetic tree with a GTR+F+I nucleotide substitution model and rooted with the best fit based on residual mean squared test statistic. It shows the geographical distribution of different SARS-CoV-2 samples across two temporally distinct lineages.

### Mutations in SARS-CoV2 peptides

CoVa reported 5 deletions and 361 point mutations analyzing Indian genomes. It also found a 1 base insertion present in a single genome. 346 of the 361 point mutations were located in peptide-encoding regions, including 224 missense and 5 nonsense mutations. All nonsense mutations were singletons, except ORF8 (E110*), which was also present only in 2 genomes. 70 of the 224 missense mutations were present in more than 1 genome. We found 28 of these mutations to be present in the global sample, 10 of which were enriched in the Indian sample (*P*_Fisher’s test_ < 0.01, adjusted for multiple testing). However, it should be noted that the Indian genomes were sequenced relatively more recently. More specifically, ~ 70 % of the Indian genomes were sequenced after 31st March, whereas only ~ 24% genomes were sequenced in the same period in the global sample. To reduce this temporal component of the observed difference, we compiled another dataset for comparison with the local sample. We also used this chance to reduce the sampling bias in the global sample, originating from a disproportionately large number of sequences from some countries. First, we selected all high quality genomes from GISAID, collected from 1st of April to 5th of May. We normalized sample size for each country, represented in this dataset, to its population as follows:

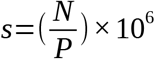

Where *s =* number of genomes sequenced per million individuals (*pmi*) from the country, *N =* number of genomes & *P =* Country’s population.

The value of *s* for India was 0.16. Therefore, we selected 0.2 pmi most recent genomes from all other countries. For countries with fewer sequences, we used all of the available data. After removing the duplicate sequences, we had a set of 250 “recent” sequences. The above 10 mutations were enriched in Indian sequences relative to global samples, even after accounting for temporal and sampling bias (**table 2**).

**Table 2.**
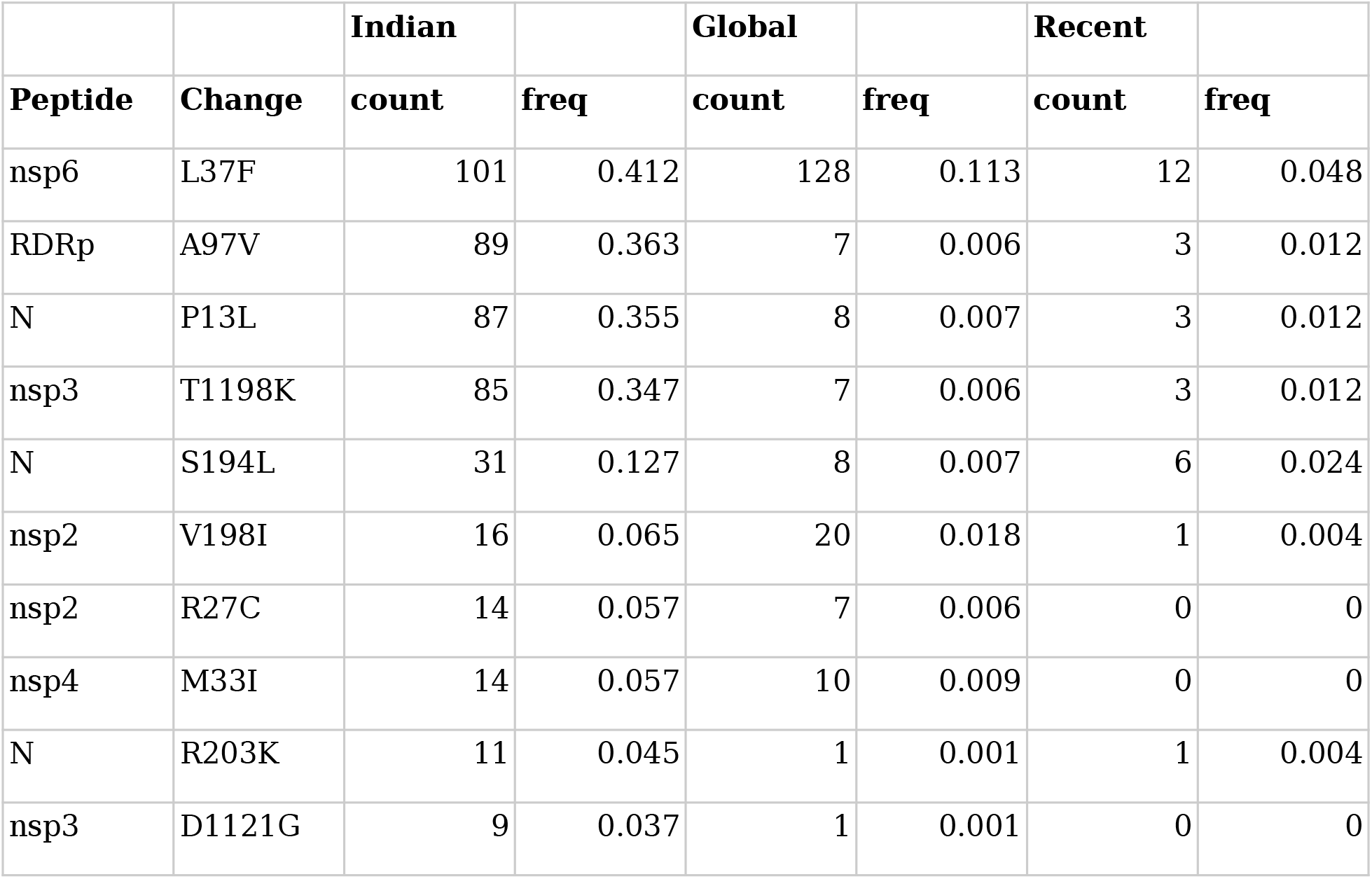
Mutations enriched among Indian sequences relative to genomes from other countries.

|  |  | **Indian** |  | **Global** |  | **Recent** |  |
| --- | --- | --- | --- | --- | --- | --- | --- |
| **Peptide** | **Change** | **count** | **freq** | **count** | **freq** | **count** | **freq** |
| nsp6 | L37F | 101 | 0.412 | 128 | 0.113 | 12 | 0.048 |
| RDRp | A97V | 89 | 0.363 | 7 | 0.006 | 3 | 0.012 |
| N | P13L | 87 | 0.355 | 8 | 0.007 | 3 | 0.012 |
| nsp3 | T1198K | 85 | 0.347 | 7 | 0.006 | 3 | 0.012 |
| N | S194L | 31 | 0.127 | 8 | 0.007 | 6 | 0.024 |
| nsp2 | V198I | 16 | 0.065 | 20 | 0.018 | 1 | 0.004 |
| nsp2 | R27C | 14 | 0.057 | 7 | 0.006 | 0 | 0 |
| nsp4 | M33I | 14 | 0.057 | 10 | 0.009 | 0 | 0 |
| N | R203K | 11 | 0.045 | 1 | 0.001 | 1 | 0.004 |
| nsp3 | D1121G | 9 | 0.037 | 1 | 0.001 | 0 | 0 |

Additionally, 7 mutations, which were present in more than 5 Indian sequences (**Sup. table 4**), were missing from the global samples. However, these are not necessarily unique to India, as we did find 3 of the 7 - nsp3 (S1197R, S697F) & nsp8 (Q198H) - in a different global dataset (not included in this study).

### Proteins under positive selection

We estimated the mode and strength of selection acting on SARS-CoV2 peptides using FUBAR over the entire phylogeny, as implemented in CoVa. FUBAR estimates gene-specific distribution of synonymous and non-synonymous rates using a Bayesian approach coupled with Markov Chain Monte Carlo (MCMC) sampling. Sites are assumed to evolve independently under a codon-based model in which rates are drawn from a prespecified discrete bivariate distribution to maximize the probability of observed alignment. FUBAR estimates these rates over the entire phylogeny and not specific branches and therefore, it is suited to detect pervasive selection. A protein or a site is considered under positive selection if its non-synonymous rate (β) is greater than the synonymous rate (α). Analysing the global dataset, we found 2 non-structural proteins - nsp2 & nsp8, and the nucleocapsid protein N to be under positive selection. Within India, we detected signs of positive selection on N, along with several other proteins - methyltransferase, ORF8, ORF3a, nsp10 & nsp11, but not on nsp2 and nsp8.

It is possible that some of these proteins are also under selection in the global dataset. However, as mentioned earlier, FUBAR detects pervasive and not episodic selection. The presence of a large no. of sequences from different parts of the world, representing independent populations, could have masked signs of positive selection on these peptides within some countries. Conversely, nsp2 and nsp8 might be under positive selection in a specific country with a large number of sequences. To test if this were indeed the case, we separated global data by countries and performed selection analysis separately (using CoVa) on 12 countries with at least 20 genomes. As suspected, we detected positive selection on these peptides in several other countries except for nsp11, which showed signs of positive selection only in India (**Figure 5**). On the other hand, nsp2 and nsp8 were actually under positive selection in many countries. β - α values for all peptides are provided in **Sup. table 5**. It should be noted that many peptides appeared to be under positive selection in a few countries, and India was not peculiar in having 6 peptides under positive selection.

**Figure 5:**
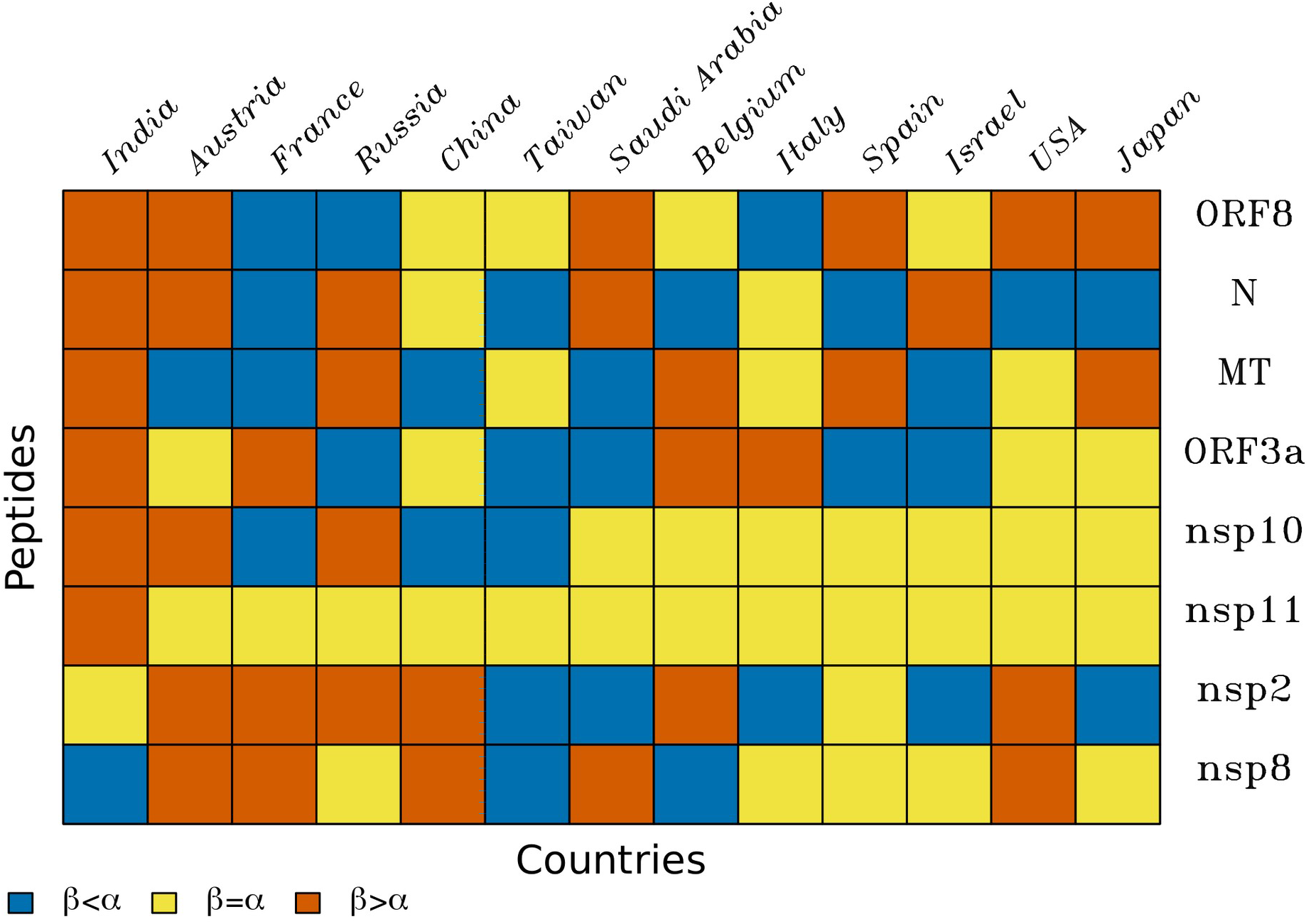
Proteins under positive selection across countries. Difference between non-synonymous substitution rate (*β*) and synonymous rate (*α*) for 8 proteins with *β - α* > 0 either in India or globally.

To test if there were any geographical correlates to the probability of a protein being under positive selection, we compared the distribution of latitudes for countries where a protein was under positive selection to that of the other countries. For nsp2, we found that the protein was under positive selection in temperate locations more often than in subtropics (**Figure 6**) (*P*_Wilcoxon rank sum_ = 0.01).

**Figure 6:**
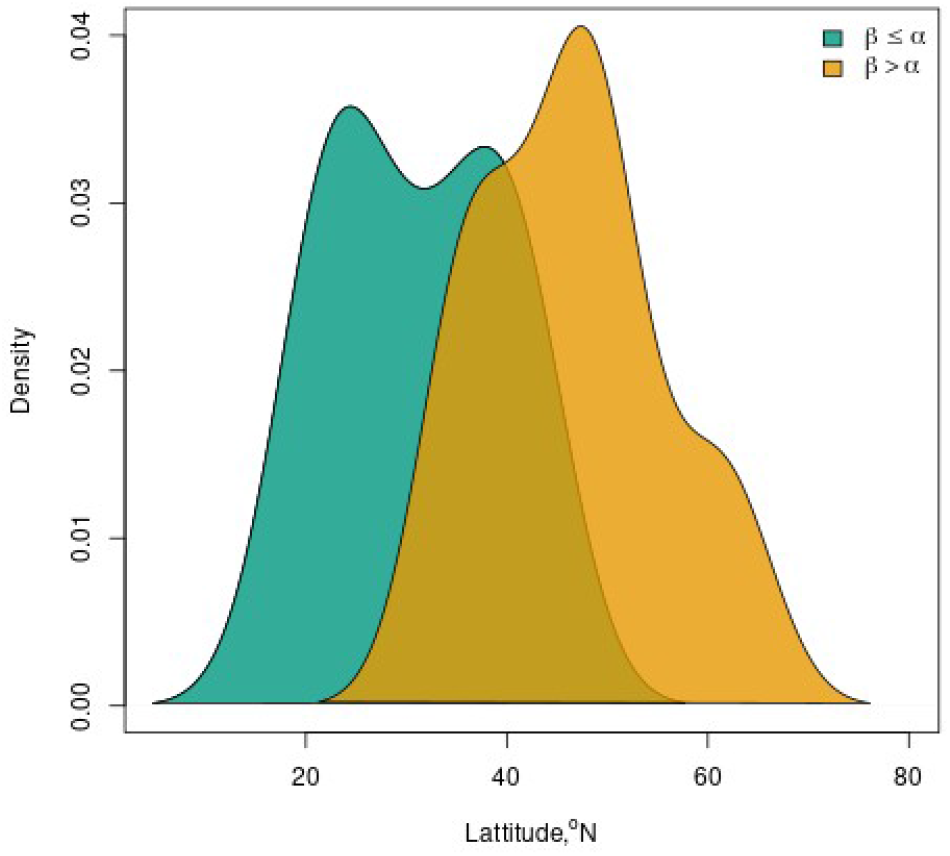
Geographical pattern of selection on nsp2. Distributions of geographic latitudes of countries with and without positive selection on nsp2.

Nsp2 is involved in viral RNA synthesis (29), and can potentially impair host cell signaling via its interaction with Prohibitins (30). An amino acid deletion (D268) was reported from France, Netherland and England, exemplifying the above trend (31). A point mutation in nsp2, T85I, was reported in sequences from the USA, and was predicted to cause structural alterations (32). We found the same mutation in 222 global sequences while only in 2 sequences from India. Nsp8 encodes an RNA polymerase which was proposed to produce primers for the canonical RDRp (nsp12) (33). Nsp11 is a small and uncharacterized 13-residue long peptide. It was shown to enhance binding of nsp10 with nsp14 (exoribonuclease) and nsp16 (methyltransferase), and thus, likely have a role in replication/transcription (34).

### Positive selection on individual sites

Notwithstanding the signs of positive selection on the proteins above, it is well known that proteins are rarely under positive selection over their entire sequence. Rather, it is more common to find a few sites under positive selection in an otherwise conserved protein. FUBAR identifies such sites based on the total posterior probability of β > α, averaged over MCMC samples, being greater than 0.9. In India, we found 27 sites as targets of positive selection distributed over 14 proteins, including the polymerase RDRp, spike protein S, and even nsp2. Only 7 of these sites could be detected in the global dataset, *viz.*, RDRp (97, 323), N (13, 194), ORF3a-57, Helicase-206 and nsp6-37.

Amino acid positions on a protein sequence are rarely the target of selection by themselves unless they solely dictate a function. We can expect groups of sites to be under selection. However, many such sites might skip detection while using a stringent and arbitrary threshold, as above. For this reason, we extracted posterior probabilities *P*_β > α_ of all the sites of these 14 proteins from 13 countries including India (**Sup. table 6**). We observed 3 distinct distributions of these values; a majority of the sites had *P*_β > α_ in the range 0.4-0.6 while a few sites had the probability > 0.7 indicating some degree of positive selection or < 0.2 suggesting purifying selection. Similar to our analysis with the whole proteins, we sought to identify sites for which the probability of positive selection was correlated with geographical parameters. We tested the strength of spearman correlation between a country’s latitude and a site’s *P*_β > α_ for 589 sites, which had *P*_β > α_ > 0.7 in at least one country. After correcting for multiple testing, we did not find a single site with a significantly non-zero correlation (*P* < 0.05). However, based on uncorrected p-values, we found 2 sites (212, 228) in nsp2 with a strong positive correlation (0.785) with latitude and the lowest *P* (0.001), supporting our observation at the protein level. Besides, multiple sites in RDRp (141, 196, 323, 668, 874), a single site in S (723) and few other sites showed negative correlation values (unadjusted *P* < 0.05), suggesting greater odds of their being positively selected at lower latitudes (**Table 3**).

**Table 3.** Sites with a strong correlation of *P*_β > α_ in a country with its latitude (|ρ| > 0.6, *P* < 0.05).

| SN | Protein | Site | rho | P | adjP |
| --- | --- | --- | --- | --- | --- |
| 1 | RDRp | 141 | -0.634 | 0.02 | 1 |
| 2 | RDRp | 323 | -0.616 | 0.025 | 1 |
| 3 | RDRp | 874 | -0.664 | 0.013 | 1 |
| 4 | S | 723 | -0.613 | 0.026 | 1 |
| 5 | nsp3 | 1198 | -0.624 | 0.023 | 1 |
| 6 | orf3a | 8 | 0.615 | 0.025 | 1 |
| 7 | orf3a | 67 | 0.64 | 0.019 | 1 |
| 8 | orf3a | 102 | 0.615 | 0.025 | 1 |
| 9 | orf3a | 222 | 0.601 | 0.03 | 1 |
| 10 | orf3a | 239 | 0.624 | 0.023 | 1 |
| 11 | N | 13 | -0.675 | 0.011 | 1 |
| 12 | N | 41 | 0.64 | 0.018 | 1 |
| 13 | N | 378 | 0.615 | 0.025 | 1 |
| 14 | nsp2 | 198 | -0.609 | 0.027 | 1 |
| 15 | nsp2 | 212 | 0.785 | 0.001 | 0.589 |
| 16 | nsp2 | 228 | 0.785 | 0.001 | 0.589 |
| 17 | nsp2 | 384 | 0.632 | 0.021 | 1 |
| 18 | nsp2 | 478 | 0.634 | 0.02 | 1 |
| 19 | nsp2 | 548 | 0.785 | 0.001 | 0.589 |
| 20 | nsp2 | 555 | 0.739 | 0.004 | 1 |

As proteins often interact in a physical space to perform a function, a group of sites under the same selection pressure can be distributed across multiple proteins. A consequence of their coupled evolution would be that if one of the co-evolving sites is under positive selection in any population, then we can expect its partner(s) to also be under positive selection in that population. With this reasoning, we clustered sites which were all under selection (P_β > α_ > 0.7) in the same set of countries. We identified 36 sites constituting 10 such clusters (**Figure 7**). Cluster 3 represented 3 sites under selection across all countries. Cluster-1, 2, 5 & 6 had sites from India common with different sets of countries. Cluster 7 was the largest with 6 sites, and it represented sites common between the USA and Taiwan. As can be seen from the figure, certain groups of proteins co-appeared in multiple clusters; nsp2 + nsp3 in 4 and S + nsp3 in 3 clusters.

**Figure 7:**
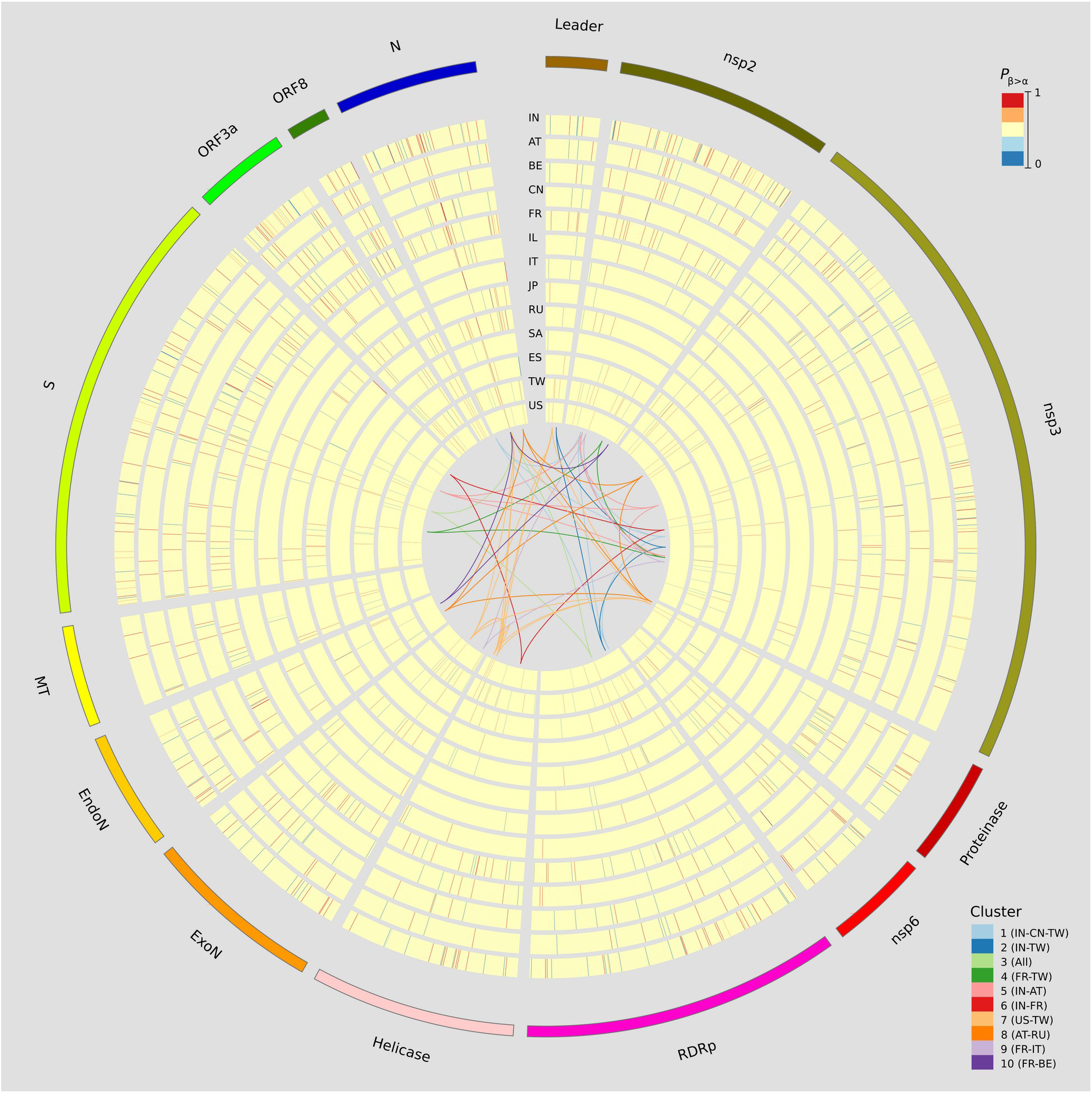
Range of selection pressure on sites of SARS-CoV-2 proteins. Majority of sites were found to be under neutral selection whereas sites under purifying or positive selection were sparse and scattered across proteins. Several sites appeared under positive selection together across a set of countries. Positively selected sites in some countries were conserved in others.

## Conclusion

Cova’s speed, accuracy and breadth of analysis makes it an ideal tool for genome analysis of SARS-CoV-2. While several other genome analysis tools (35–37) have been developed for this virus, CoVa accomplishes all of these analyses in a short time and with minimum computational resources. As opposed to other web-based tools, CoVa can be run locally and, being a command-line utility, has the added advantage that it can be easily integrated into a larger workflow for bioinformatics research. The present capabilities of CoVa allowed us to identify subtypes of SARS-CoV-2, both in India and across the world. Using CoVa, we could detect signs of positive selection across multiple proteins and individual sites. Interesting patterns emerged from this study across geographies. Some of these might be explained by environmental factors, like ambient temperature, relative humidity etc, others might require epidemiological explanations, demanding more detailed analysis. Future updates to CoVa will include more sophisticated phylogenetic analysis methods and mutation mapping to available structure data.

## Ethical Approval

The use of patient-derived SARS-CoV-2 genome sequences from the GISAID database was approved by the NCBS Institutional Human Ethics Committee vide letter NCBS/IEC-19/002.

## Acknowledgement

We gratefully acknowledge the authors, originating and submitting laboratories of the sequences from GISAID’s EpiFlu™ Database on which this research is based. The complete list is available in **supplementary tables 7-9**.

## Availability

Cova is available to download from https://github.com/A-Farhan/cova. The instructions to install and run CoVa, along with example datasets are also included

